# Factors affecting genetic connectivity and diversity of the island night lizard

**DOI:** 10.1101/060038

**Authors:** Stephen E. Rice, Rulon W. Clark

## Abstract

Habitat loss and fragmentation is one of the most severe threats to global biodiversity. Because human development often fragments natural areas into habitat “islands”, studies which characterize the genetic structure of species isolated on oceanic islands may provide insight into the management of anthropogenic habitat islands. The San Clemente Island night lizard, *Xantusia riversiana reticulata*, is endemic to two California Channel Islands, each with a history of anthropogenic disturbance. We genotyped 917 individuals from San Clemente Island and Santa Barbara Island at 23 microsatellite loci to quantify population structure and identify natural and anthropogenic landscape features affecting intra-island connectivity. We found significant, but shallow, population structure on each island with sites < 400 m apart identified as distinct genepools. Landscape genetic analyses identified conductive habitat as California boxthorn and prickly pear cactus on both islands. Landscape features which decreased connectivity were unique to each island and included natural and human-mediated features. These results can inform management plans on each island by identifying habitat targets for mitigation and restoration efforts designed to improve connectivity. Our results highlight the need for considering fine-scale features correlated with contemporary and historical patterns of fragmentation, especially in small and isolated habitats on the mainland that may be analogous to oceanic islands.

## Introduction

Habitat fragmentation is a leading cause of global biodiversity loss. Fragmentation can decrease metapopulation connectivity, increase genetic divergence, and increase extinction risk (e.g. Frankham 2005; Sumner 2005; Vandergast et al. 2016). Fragmented and isolated habitat patches often form “habitat islands”, patches of suitable habitat embedded within an inhospitable matrix, such as a nature preserve surrounded by human housing developments (e.g. Hokit et al. 1999; Driscoll 2004; Wang et al. 2009). When this occurs, the ecological effects of fragmentation can be further exacerbated by management practices, as isolated fragments may be managed independently by different agencies, which may result in variable outcomes in terms of conservation goals, even when different fragments share common species and habitats. Thus, one goal of modern conservation is to identify generalizable approaches which can be readily implemented across administrative boundaries to improve the management of fragmented and isolated habitat patches. This approach requires examining genetic connectivity and diversity both within and between fragments, in order to distinguish features shared in common among fragments that can impede or enhance connectivity, as well as features that may be specific to each fragment. Although these types of comparisons are often made in the context of actual oceanic islands, they are less commonly applied to habitat islands. Here, we conduct a management case study of a species distributed across two oceanic islands managed by different agencies. We believe this and other case studies can serve as valuable exemplars for conservation approaches that could also be applied to species distributed across habitat islands.

The California Channel Islands are small oceanic islands off Southern California’s coast which share the Mediterranean climate of the mainland and have experienced habitat degradation and fragmentation due to anthropogenic activities (e.g. Fellers and Drost 1991; Fellers et al. 1998; Sturgeon 2000). Different islands are managed by different government agencies, although they share several species of conservation concern in common. Thus characterizing the contemporary genetic patterns of Channel Island endemics will also provide critical data for monitoring and management directed toward island endemics (Schwartz et al. 2007; reviewed in Stetz et al. 2011) and serve as a case study for habitat islands.

Reptiles are an excellent group for studies of fragmentation due to their intermediate dispersal distances, relatively high population densities, plasticity of responses, thermal physiology, and potential vulnerability to habitat modifications (e.g. Hokit et al. 1999; Jordan and Snell 2008; Wang et al. 2009; Tseng et al. 2015). One of the few reptiles endemic to the Channel Islands is the island night lizard, *Xantusia riversiana* (Cope 1883). This species was delisted from the Endangered Species Act in 2014, contingent on continued monitoring (United States Fish and Wildlife Service (USFWS) 2014). Two subspecies are recognized (Smith 1946), *X. r. reticulata*, the San Clemente Island night lizard, endemic to San Clemente Island and Santa Barbara Island (Figure 1) and *X. r. riversiana* endemic to San Nicolas Island. These subspecies differ in both life history traits and habitat use (Fellers and Drost 1991; Fellers et al. 1998). On San Clemente Island lizards are found primarily in habitats dominated by California boxthorn (*Lycium californicum*) and prickly pear cactus (*Opuntia littoralis*), and avoid deep canyons, sand dunes, and canyon woodland (Mautz 1993). On Santa Barbara Island, lizards are found primarily in habitats dominated by California boxthorn and prickly pear cactus, and tend to avoid grasslands (Fellers and Drost 1991). In prime habitat *X. r. reticulata* may reach densities in excess of 3,200 individuals/ha (Fellers and Drost 1991; Mautz 1993).

**Figure 1.**
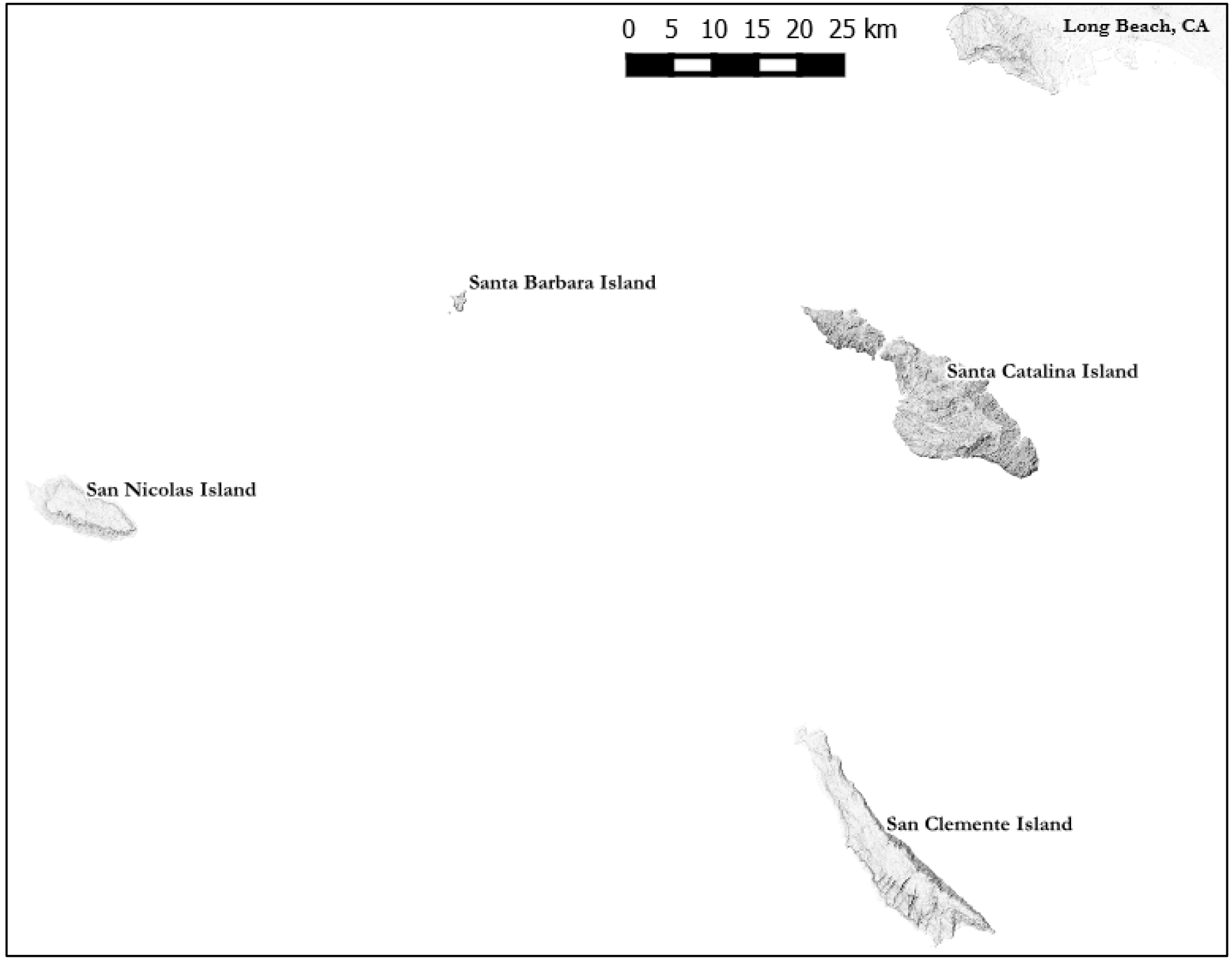
Location of Channel Islands inhabited by the island night lizard. The island night lizard is endemic to 3 Channel Islands: San Clemente Island, Santa Barbara Island, and San Nicolas Island.

Although the island night lizard can reach very high population densities in prime habitat, this long-lived species may be vulnerable to habitat fragmentation and disturbance due to a combination of life history traits and very low dispersal distances (Fellers and Drost 1991; Mautz 1993). As part of an ongoing post-delisting monitoring effort, we characterized the genetic structure and connectivity of *X. r. riversiana* across its entire geographic range, sampling sites spread across both San Clemente and Santa Barbara Islands. We also characterized habitat, anthropogenic structures, and other geographic features that may serve as potential barriers to population connectivity. Based on large population sizes and near-continuous distribution of viable habitat, we hypothesized that population genetic structure would be minimal on each island but influenced by roads and other anthropogenic barriers, as well as non-preferred habitat types. We also hypothesized that detected patterns of genetic differentiation would be primarily correlated with Euclidean distance, due to limited dispersal (Fellers and Drost 1991; Mautz 1993).

## Materials and Methods

### Sample Collection and Genotyping

We collected samples on San Clemente Island (Figure 2) from February through November of 2013 and Santa Barbara Island (Figure 3) from May through September 2015. Island night lizards were captured by turning cover items and setting live traps along well-worn trails. When multiple lizards were encountered, we made attempts to capture and sample all lizards. Target sample size for each collection site was a minimum of 30 individuals (Hale et al. 2012). Samples were also collected opportunistically from encountered individuals. Captured lizards were processed following the United States Geological Survey herpetofaunal monitoring protocols (Fisher et al. 2008) and GPS coordinates were recorded with an estimated accuracy of 5 m. Tissue samples consisted of toes that were clipped at the distal knuckle, which also served as a unique 4-digit identification, and a 10 mm segment from the tail tip. Tissues were preserved in 95% ethanol and stored at –20°C.

**Figure 2.**
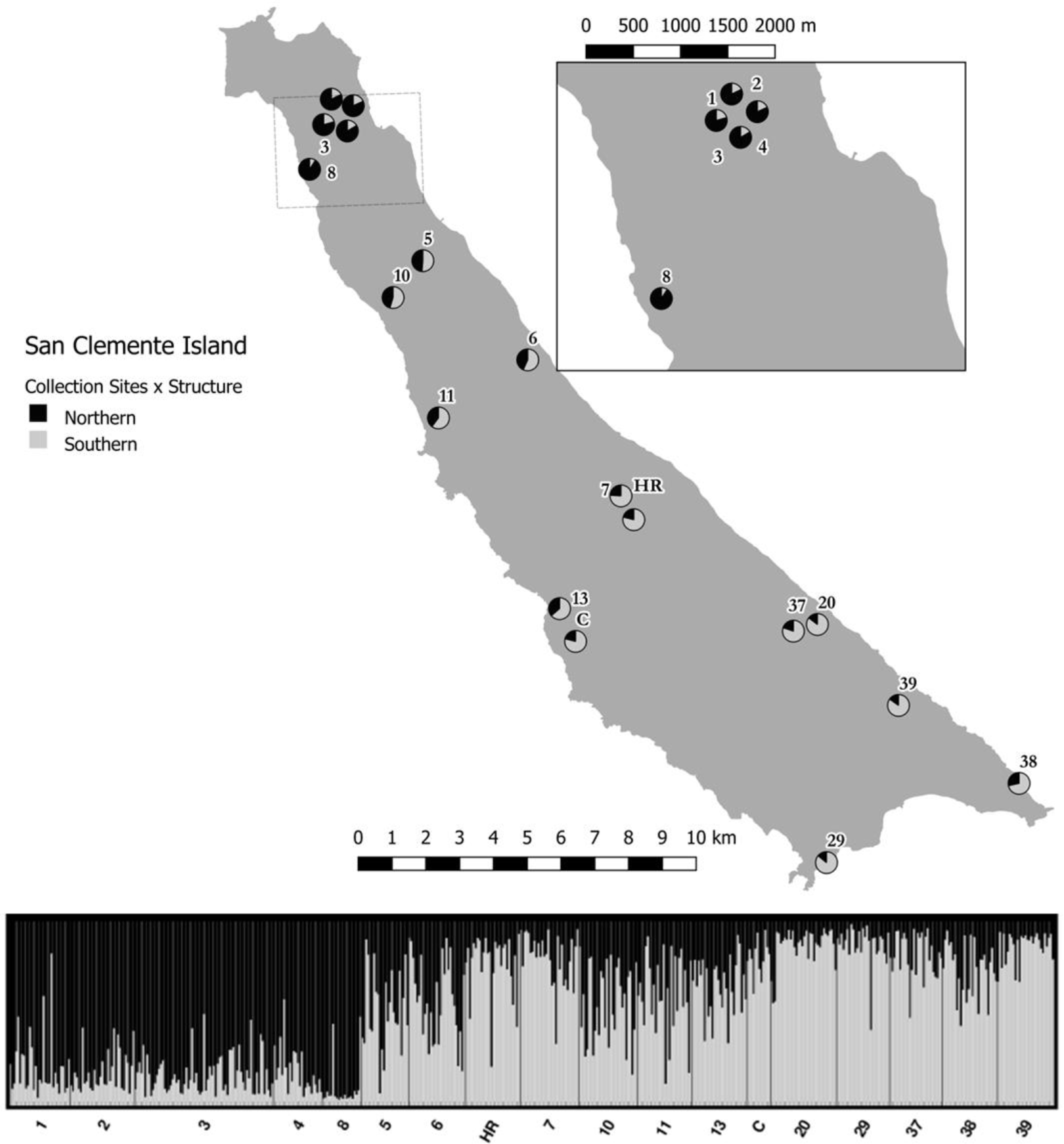
San Clemente Island collection sites with coancestry proportions. All collection sites on San Clemente Island are shown. Pie charts represent the site-level coancestry proportions for the STRUCTURE solution K=2 (barplot).

**Figure 3.**
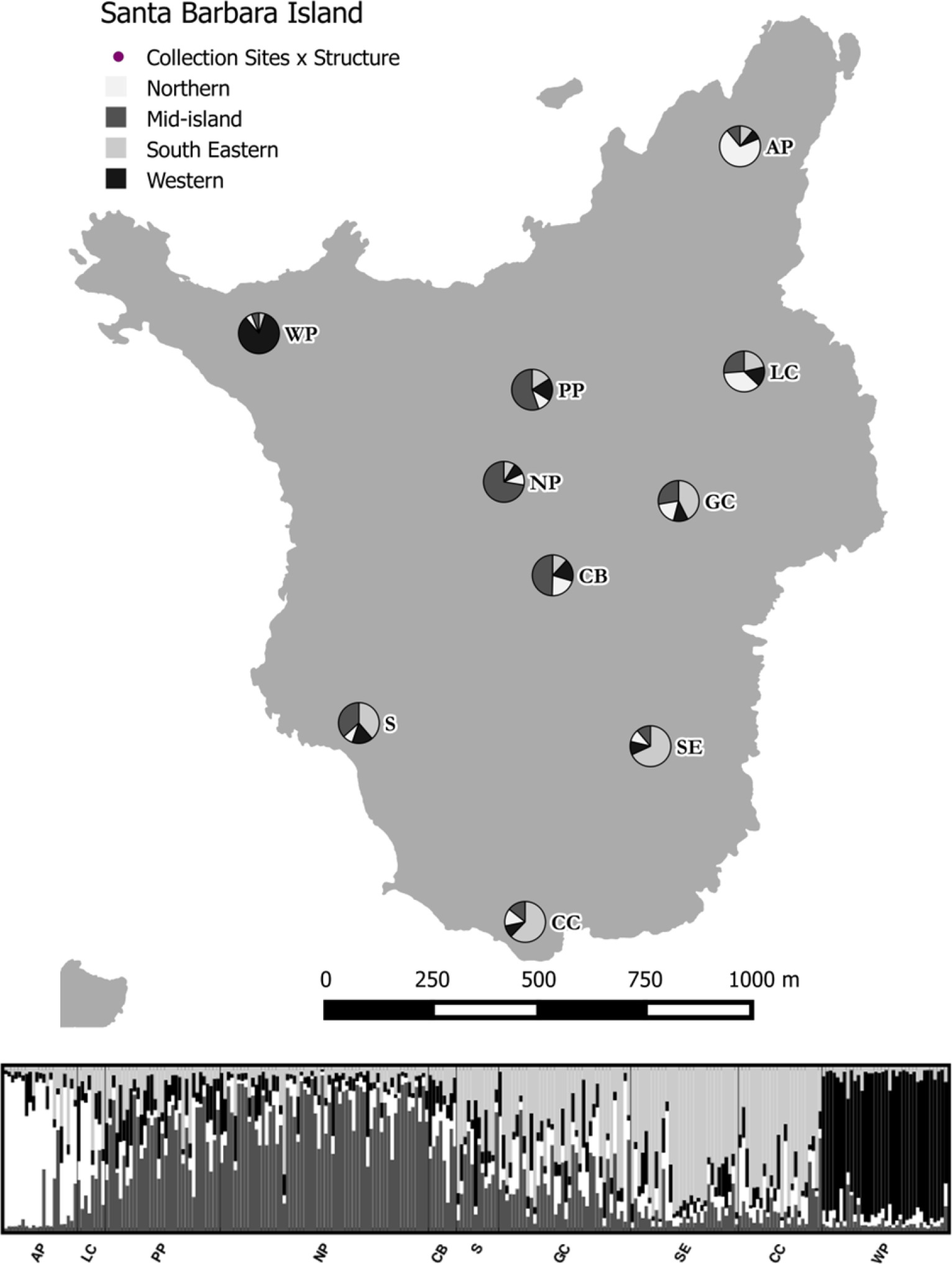
Santa Barbara Island collection sites with coancestry proportions. All collection sites on Santa Barbara Island are shown. Pie charts represent the site-level coancestry proportions for the STRUCTURE solution K=4 (barplot).

We extracted DNA with a standard salt digestion and genotyped individuals at 23 polymorphic microsatellite loci arranged into 8 reactions (Rice et al. 2016; Rice 2017 Appendix A, B). We scored alleles twice with Genemapper for consistency; loci missing data or with inconsistent allele calls were reamplified and rescored to generate a consensus genotype. We assessed per-allele error rates (Pompanon et al. 2005) by extracting and genotyping 35 individuals on San Clemente Island and 16 individuals on Santa Barbara Island a second time.

### Quality Control and Summary Statistics

We removed individuals from analyses if amplification was successful at fewer than 16 loci. We removed first-order relatives (full siblings, parent-offspring) from data sets for each island to prevent biases in detected population structure. We inferred pairwise relationships for samples within each collection site when at least 2 of 3 methods were concordant. The methods we chose for inference were Colony (Jones and Wang 2010), Cervus (Kalinowsi et al. 2007), and the DyadML estimator as calculated in Coancestry (Wang 2011). Individuals were removed from each data set to retain the largest sample size. Final data sets with relatives removed were used in all further analyses.

We evaluated the quality of each locus for each island at the level of collection sites following Selkoe and Toonen (2006). Loci were removed if amplification occurred in fewer than 75% of individuals, when in linkage disequilibrium (LD) with another locus, when null alleles occurred in a majority of sampled sites, or when the assumption of neutrality was violated. We examined loci for LD and conformance to Hardy-Weinberg and random mating proportions (HWE) in Genepop (Rousset 2008) with p-values corrected for multiple comparisons through sequential Bonferroni testing (Holm 1979). We assessed the presence of null alleles with Microchecker (Van Oosterhout et al. 2004). We examined the putative neutrality of loci with Lositan (Beaumont and Nichols 1996; Antao et al. 2008) under the infinite alleles model and neutral Fst estimation. We used the package diveRsity (Keenan et al. 2013) in the R statistical environment and Genepop to generate summary statistics for each collection site and each island (Table 1).

**Table 1.** Summary statistics for each island and collection site. Collection sites are listed under their respective islands with the total samples for that location (N) and those in the reduced data set (Nrr). Summary statistics were: rarified allelic richness for all collection sites (Ar), rarified allelic richness for only those sites with ≥ 20 individuals in the reduced data set (Ar (≥20)), expected heterozygosity (He), observed heterozygosity (Ho), HWE probabilities (HWE) with probabilities <0.0001 set to 0.0001 at the locus level. Island-level comparisons were generated from the reduced data sets, analyzed at the 18 shared loci. The inbreeding coefficient (Fis) was calculated by allele identity within GENEPOP overall and for each site. NA denotes analysis not applicable.

| Site | N | Nrr | Ar | Ar ($\geq 20$) | He | Ho | HWE | Fis |
| --- | --- | --- | --- | --- | --- | --- | --- | --- |
| San Clemente Island | 609 | 534 | 21.47 | NA | 0.89 | 0.84 | $< 0.0001$ | 0.0459 |
| 1 | 33 | 31 | 8.98 | 10.35 | 0.86 | 0.87 | 0.8981 | 0.0137 |
| 2 | 39 | 33 | 8.78 | 10.08 | 0.86 | 0.86 | 0.2835 | 0.0148 |
| 3 | 86 | 71 | 9.31 | 11.0 | 0.86 | 0.87 | 0.1168 | 0.0027 |
| 4 | 30 | 25 | 8.53 | 9.77 | 0.86 | 0.87 | 0.6173 | 0.0019 |
| 5 | 28 | 24 | 8.59 | 9.92 | 0.86 | 0.88 | 0.3311 | 0.0018 |
| 6 | 31 | 29 | 8.24 | 9.50 | 0.84 | 0.86 | 0.4165 | 0.0044 |
| HR | 36 | 28 | 8.07 | 9.25 | 0.84 | 0.86 | 0.3063 | -0.0008 |
| 7 | 32 | 30 | 8.21 | 9.49 | 0.84 | 0.83 | 0.2200 | 0.0317 |
| 8 | 27 | 20 | 5.46 | 6.28 | 0.69 | 0.62 | 0.0005 | 0.1157 |
| 10 | 30 | 30 | 9.20 | 10.80 | 0.86 | 0.87 | 0.1464 | -0.0022 |
| 11 | 31 | 28 | 8.55 | 9.94 | 0.85 | 0.82 | 0.0089 | 0.0508 |
| 13 | 29 | 28 | 9.07 | 10.69 | 0.87 | 0.88 | 0.1188 | 0.0089 |
| C | 13 | 12 | 7.34 | NA | 0.82 | 0.86 | 0.4647 | -0.0056 |
| 20 | 36 | 34 | 8.27 | 9.57 | 0.84 | 0.85 | 0.0838 | 0.0023 |
| 29 | 29 | 27 | 8.17 | 9.28 | 0.85 | 0.86 | 0.0488 | 0.0062 |
| 37 | 34 | 27 | 8.26 | 9.63 | 0.84 | 0.83 | 0.3470 | 0.0319 |
| 38 | 30 | 28 | 9.21 | 10.83 | 0.87 | 0.86 | 0.9637 | 0.0275 |
| 39 | 31 | 29 | 8.31 | 9.65 | 0.84 | 0.85 | 0.2572 | 0.0097 |
| Santa Barbara Island | 312 | 271 | 13.82 | NA | 0.86 | 0.82 | $< 0.0001$ | 0.0473 |
| AP | 28 | 21 | 5.85 | 7.20 | 0.80 | 0.80 | 0.2612 | 0.0292 |
| CB | 8 | 8 | 5.62 | NA | 0.79 | 0.83 | 0.8888 | -0.0051 |
| CC | 31 | 24 | 6.44 | 8.18 | 0.83 | 0.83 | 0.3197 | 0.0086 |
| GC | 39 | 38 | 6.65 | 8.65 | 0.83 | 0.84 | 0.6265 | 0.0134 |
| LC | 9 | 8 | 5.97 | NA | 0.81 | 0.89 | 0.9830 | -0.0208 |
| NP | 69 | 60 | 6.49 | 8.42 | 0.82 | 0.82 | 0.0227 | 0.0134 |
| PP | 37 | 33 | 6.28 | 7.87 | 0.82 | 0.81 | 0.1433 | 0.0251 |
| S | 13 | 12 | 5.92 | NA | 0.79 | 0.77 | 0.1239 | 0.0581 |
| SE | 35 | 31 | 6.56 | 8.36 | 0.83 | 0.85 | 0.6892 | 0.0026 |
| WP | 43 | 36 | 5.67 | 7.17 | 0.78 | 0.76 | 0.1836 | 0.0453 |
| Aggregate LC+GC | 48 | 46 | 6.77 | 8.88 | 0.84 | 0.84 | 0.3103 | 0.0089 |

### Population Structure

We delineated genepools on each island following Waples and Gaggiotti (2006). We conducted an exact test for genetic differentiation for all pairs of collection sites in Genepop. We combined probabilities with Fisher’s method (Fisher 1932) with 0.0001 as the lower bound for individual test probabilities; combined probabilities were corrected using sequential Bonferroni testing. Collection sites were grouped if no significant difference was found and the procedure was repeated until all pairwise comparisons were significant.

We estimated genetic differentiation between collection sites with ≥ 20 individuals with Weir and Cockerham’s (1984) estimate of Fst with 95% bias corrected confidence intervals generated by 1,000 bootstraps over individuals in diveRsity. We defined differentiation as significant when the confidence interval did not overlap 0.

We used the program STRUCTURE (Pritchard et al. 2000; Falush et al. 2003) to estimate the number of populations on each island (K) using data from all collection sites. We used values of K from 1 to 18 for San Clemente Island and 1 to 10 for Santa Barbara Island. Analyses for both island consisted of 20 runs with a 500,000 burn-in period and 1,000,000 MCMC iterations under the admixture model with uncorrelated and correlated allele frequency models (Pritchard et al. 2010). We used the Evanno et al. (2005) method as implemented in Structure Harvester (Earl and vonHoldt 2012) to estimate the number of populations. We generated ancestry matrices with the greedy algorithm in the program Clumpp (Jakobsson and Rosenberg 2007) to produce barplots with the program Distruct (Rosenberg 2004) for the best supported value of K for each island.

### Isolation by Distance

We examined patterns of isolation by distance (IBD, Wright 1943) at the level of collection sites for each island. We used non-parametric rank-based multiple regression on distance matrices (MRDM, Lichstein 2007) using the function *MRM* in the R package Ecodist (Goslee and Urban 2007). As an extension of the Mantel test framework, results of MRDM are comparable to Mantel results with a single predictor matrix. Analyses were ran with 10,000 permutations, pairwise Fst as the response variable, and pairwise Euclidean distance as the predictor.

### Landscape Features Affecting Connectivity

We used the isolation by resistance framework (IBR, McRae 2006) as implemented in the program Circuitscape (McRae and Beier 2007) to model conductance and resistance of landscape features on each island. Resistance and conductance were modeled separately to clearly delineate landscape features with positive and negative correlations with pairwise genetic distances. Raster cell values for each feature ranged from 2 (low), 50 (moderate), and 100 (high). These values were chosen to identify candidate landscape features and attribute a coarse value to each for modelling purposes only and were not chosen to represent the true and unknown values of landscape features. Collection sites were represented as focal regions composed of the minimum-spanning convex hulls generated from capture coordinates.

We used non-parametric MRDM with 10,000 permutations to assess the categorical features for each island. Models ranged from single features to full models of all statistically supported features. Single features entered the model-building framework if their inclusion was significant and the coefficient of determination (R^2^) was greater than the null model. The null model consisted of bounded Euclidean distance wherein all terrestrial raster cells had a value of 1. In full models, raster cells had either the sum of resistance values for all features within the cell or 1 when no focal feature was present.

We used a consensus approach for model selection which considered the ranked-order of best models across MRDM, corrected Akaike information criterion (AlCc), and marginal 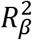 (Edwards et al. 2008; Van Strien et al. 2012). We used the *lmer* function in the R package lme4 (Bates et al. 2015) to construct maximum-likelihood population effects (MLPE) models (Clarke et al. 2002) fit with residual maximum likelihood (REML) estimation with collection site as a random factor. We used the *KRmodcomp* function in the R package pbkrtest (Halekoh and Højsgaard 2014) to calculate the marginal 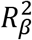 from the MLPE models. We calculated AICc with the R package MUMIN (Bartoń 2016) for MLPE models fit without REML estimation, as variables which contributed the fixed effects differed between models (reviewed in Van Strien et al. 2012).

Analyses of San Clemente Island focused on three categorical GIS layers: vegetation type (RECON, Inc 2012), roadways type (Naval Facilities and Engineering Command South West (NAVFACSW), date unknown), and canyon length (NAVFACSW, date unknown). Vegetation was parsed into 11 classes based on least-cost transect analysis groupings. Roadways were divided into two classifications: paved roads, which served as primary traffic conduits along the spine of the island and within more populated areas, and secondary roads, which were generally unpaved access roads. Canyons were divided into three length classes of small (0-499 ft), medium (500-999 ft), and large (≥ 1,000 ft). We applied a 5 m buffer to roadways and canyons to eliminate breaks in the linear features when rasterized and produced maps with a 100 m resolution

Analyses of Santa Barbara Island focused on three layers: vegetation type (National Park Service (NPS), 2010), presence of western seagull (*Larus occidentalis*) nesting colonies, and presence of a potential predator, barn owls (*Tyto alba).* There are no roads on Santa Barbara Island. Vegetation was classified into 14 groups based on dominant vegetation alliances; due to observations of woolly seablite (*Suaeda taxifolia*) dominated habitat transitioning seasonally into crystalline iceplant (*Mesembryanthemum crystallinum*) dominated habitat (pers. comm., Rodriguez 2016) we grouped these alliances into a single class. Western seagull nesting colony presence was included by digitizing seabird nesting locations from 2015 (NPS 2015). Potential barn owl presence was included by extracting and digitizing home ranges from Figure 1 of Thomsen et al. (2014). Resolution on vegetation data was listed as 1 ft (NPS 2010); however raster maps were generated with a cell size of 5 m for all layers due to the uncertainty in GPS coordinates.

## Results

### Quality Control

We sampled 605 island night lizards from 18 collection sites on San Clemente Island and 312 island night lizards from 7 collection sites and 3 smaller opportunistic areas on Santa Barbara Island. We removed 18 individuals from the San Clemente Island data set due to missing data; no individuals were removed from the Santa Barbara Island data set for this reason. We identified 75 first-order relationships among 114 individuals on San Clemente Island and 65 first-order relationships among 97 individuals on Santa Barbara Island. We removed 53 individuals from the San Clemente Island data set and 41 individuals from the Santa Barbara Island data set due to these relationships. These removals resulted in sample sizes of 534 individuals from San Clemente Island and 271 from Santa Barbara Island.

Final data sets consisted of 20 loci for San Clemente Island and 21 loci for Santa Barbara Island. A single locus was removed due to null alleles in 12 of 18 collection sites on San Clemente Island; no loci were removed from the Santa Barbara Island data set due to null alleles. The program Lositan identified 2 loci as under selection for San Clemente Island and 2 different loci as under selection for Santa Barbara Island. These loci were removed from further analyses within each respective data set. The per-allele error rates associated with genotyping were 0.062% for San Clemente Island and 0.272% for Santa Barbara Island. Rates of non-robust amplification were 7.14% for San Clemente Island and 3.80% for Santa Barbara Island.

### Summary Statistics

Sample sizes for San Clemente Island collection sites ranged from 12-70 individuals. Rarefied allelic richness for collection sites with ≥ 20 samples ranged from 6.28 to 11.0. Observed heterozygosity ranged from 0.62 to 0.88 whereas expected heterozygosity ranged from 0.69 to 0.87. Collection site 8, which is a coastal site bordered by ocean and active sand dunes on all but one side was the only site which deviated from HWE and also had the lowest allelic richness, observed heterozygosity, and expected heterozygosity (Table 1).

Sample sizes for Santa Barbara Island collection sites ranged from 21-60 individuals and 8-12 individuals for three opportunistic collection areas. Rarified allelic richness for the 7 main collection sites ranged from 7.17 to 8.65. Observed heterozygosity ranged from 0.76 to 0.89. Expected heterozygosity ranged similarly from 0.78 to 0.83. No significant departures from HWE were detected (Table 1).

### Population Structure

All collection sites on San Clemente Island were identified as distinct genepools with combined p-values from functionally 0 to 1.02 × 10^−5^. Pairwise Fst values between collection sites revealed subtle but significant structure on San Clemente Island (Table S1, supplemental) with values ranging from 0.0022 to 0.1278. The largest values for Fst involved collection site 8. The best supported clustering solution found by STRUCTURE for San Clemente Island was K=2 (Figure 2).

All collection sites on Santa Barbara Island, except two, were distinct genepools with p-values ranging from functionally 0 to 1.71 × 10^−7^ . Summary statistics for the grouped sites (LC and GC) are presented independently and together in Table 1. Since LC was an opportunistic site with few samples, LC and GC were not aggregated for further analyses. On Santa Barbara Island pairwise Fst also showed subtle but significant structure (Table S2, supplemental) with values ranging from 0.0199 to 0.0590. Structure analyses with correlated and uncorrelated allele frequencies converged on solutions of K=4 and K=7. We chose the solution of K=4 (Figure 3) as the K=7 solution displayed clusters with minimal contributions.

### Factors Positively Affecting Connectivity

On San Clemente Island, conductance models (Table 2) that included California boxthorn (boxthorn) and prickly pear cactus (prickly pear) had the strongest support across methods. Four models were equivalent in significance (p=0.0001, R^2^ range: 0.6433-0.6602). Ranking these models by AICc led to equivalence between two models that differed only by the inclusion of grasslands as low conductance; the model with the lowest AlCc (-957.56) and highest 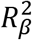 value (0.7257) contained both grasslands and small canyons as low conductance (Figure 4).

**Figure 4.**
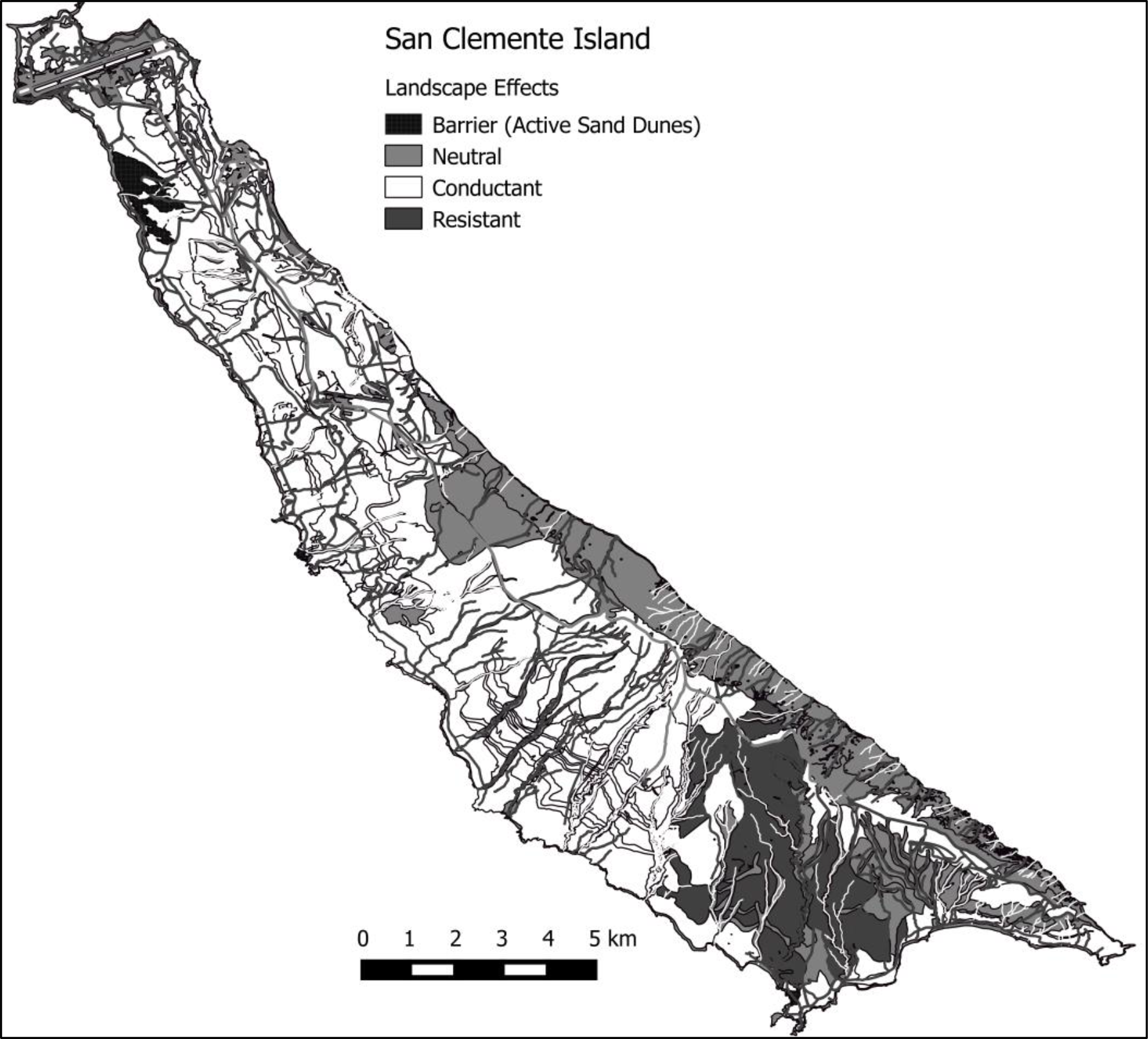
San Clemente Island landscape effects. The landscape-level features correlated with genetic distance incorporated as a composite image. Active sand dunes were identified as a barrier (black). Landscape features in gray were neutral with respect to connectivity. Conductant features (California boxthorn, prickly pear cactus, grasslands, and canyons < 500ft in length) are shown in white. Resistant features (cholla phase maritime desert scrub secondary roadways, canyons > 500ft in length) are shown in dark gray.

On Santa Barbara Island, conductance models (Table 3) which included boxthorn and prickly pear were also the best supported across methods. Eight models had AlCc values within 2 units of the best model (AICc range -151.976 to -153.318). The R^2^ for conductance models ranged from 0.8596 to 0.8877. The model with the greatest R^2^ and 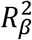 (0.8718) had both boxthorn and prickly pear with high conductance and giant coreopsis (*Leptosyne gigantean*) as low conductance (p=0.0008; Figure 5). Statistical methods did not show congruence among model rankings after the best supported model.

**Figure 5.**
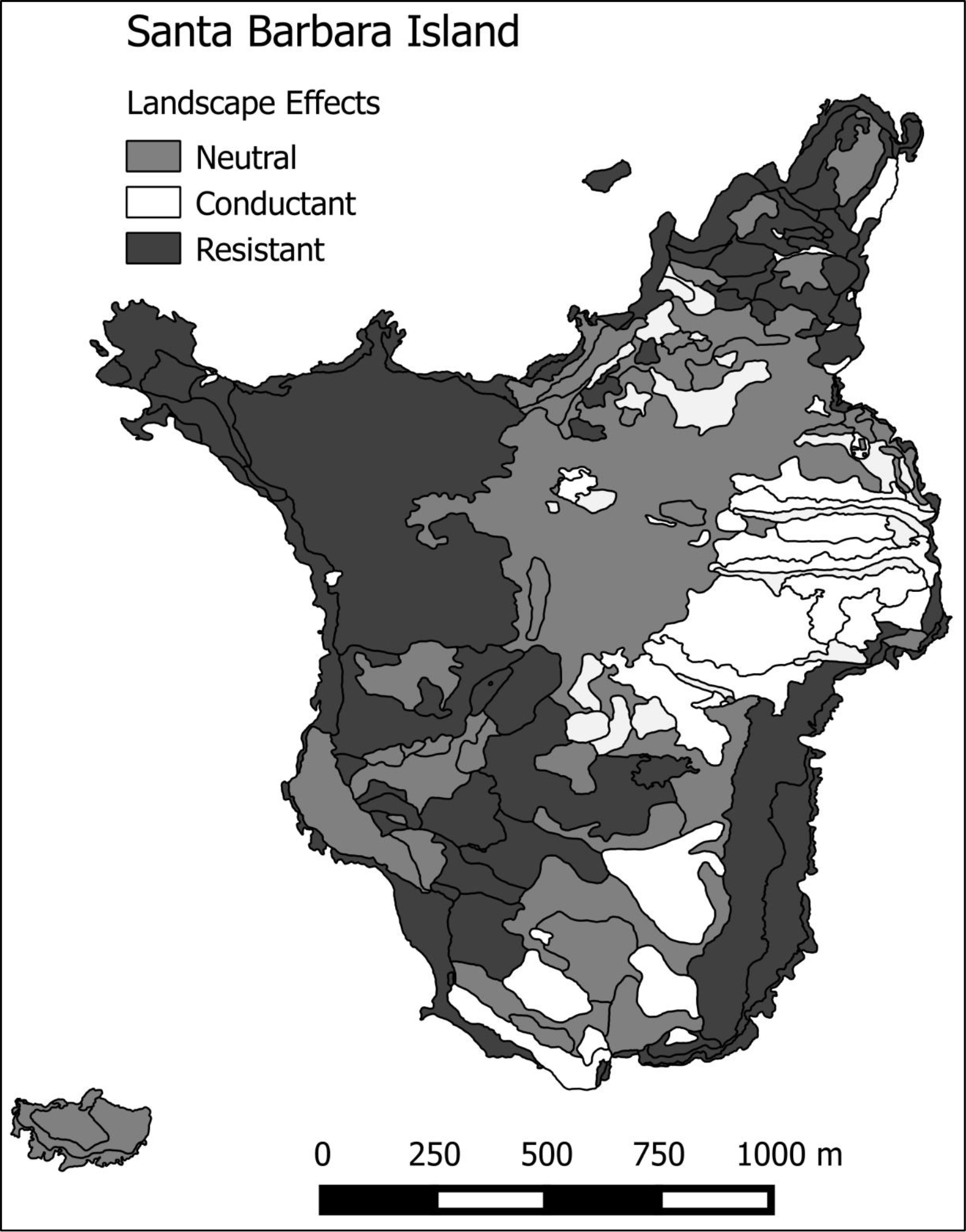
Santa Barbara Island landscape effects. The landscape-level features correlated with genetic distance incorporated as a composite image. Landscape features in gray were neutral with respect to connectivity. Conductant features (California boxthorn, prickly pear cactus, and giant coreopsis) are shown in white. Resistant habitat (woolly seablite with crystalline iceplant, barren ground, and common fiddleneck) are shown in dark gray.

### Factors Negatively Affecting Connectivity

We detected significant patterns of IBD between collection sites with ≥ 20 samples on both San Clemente Island (p=0.001, R^2^ =0.2060; Figure S3, supplemental) and Santa Barbara Island (p=0.0174, R^2^=0.4258, Figure S4, supplemental). On San Clemente Island, initial analyses resulted in no detection of IBD (p=0.145, R^2^=0.0249).We attributed this to the non-linear and non-monotonic relationship between Fst and Euclidean distance resulting from the strong signal of genetic differentiation between collection site 8 and all other sites. Removal of this site as an outlier resulted in a significant finding of IBD among the remaining collection sites.

IBR analyses of San Clemente Island were also sensitive to the inclusion of collection site 8. We evaluated 4 models to determine if the observed patterns could be attributed to collection site 8 as an outlier or active sand dunes acting as a barrier. The best supported model placed active sand dunes as a complete barrier (p=0.0005,R^2^=0.2602, Figure S3, supplemental), followed by a distance-only model with collection site 8 removed (p=0.0020, R^2^=0.1121), collection site 8 removed and active sand dunes as a complete barrier (p=0.0096, R^2^=0.0848), and a bounded Euclidean distance (IBR null) model with all collection sites (p=0.0149, R^2^=0.0681, Figure S3, supplemental).

On San Clemente Island, resistance models (Table 2) with cholla phase maritime desert scrub (cholla), canyons > 500 ft, and secondary roads were best supported across methods. The best models contained the same features, preventing model selection by AICc (AICc: -972.94 to -974.47). In the top performing models cholla, medium canyons, and secondary roadways were moderate resistance whereas large canyons were moderate to high resistance. The model best supported by MRDM (p=0.0001, R^2^=0.5125) and 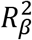 (0.8606) contained cholla, medium canyons, and secondary roads with moderate resistance and large canyons with high resistance (Figure 4).

The IBR null model on Santa Barbara Island was significant (p=0.0214,R^2^=0.4804; Figure S4, supplemental). Resistance models which included woolly seablite (seablite) and crystalline iceplant (iceplant) as moderately resistant were the best supported models (Table 3). Seven models had equivalent R^2^ (0.8582) and highly significant p-values (p < 0.001). Four of these models had AICc values of -149.043 to =150.066 and differed only in the resistance value assigned to barren ground and the inclusion of needle goldfields (*Lasthenia gracilis*). The model with the greatest 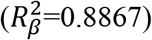 and within the AICc range contained seablite and iceplant as low resistance with barren ground as moderate 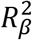 however, this model had a lower R^2^value (0.6919). We chose the model in which combined seablite and iceplant with barren ground and common fiddleneck had moderate resistance (Figure 5) as increased resistance and additional habitat types failed to greatly improve the model (Table 3).

**Table 2.**
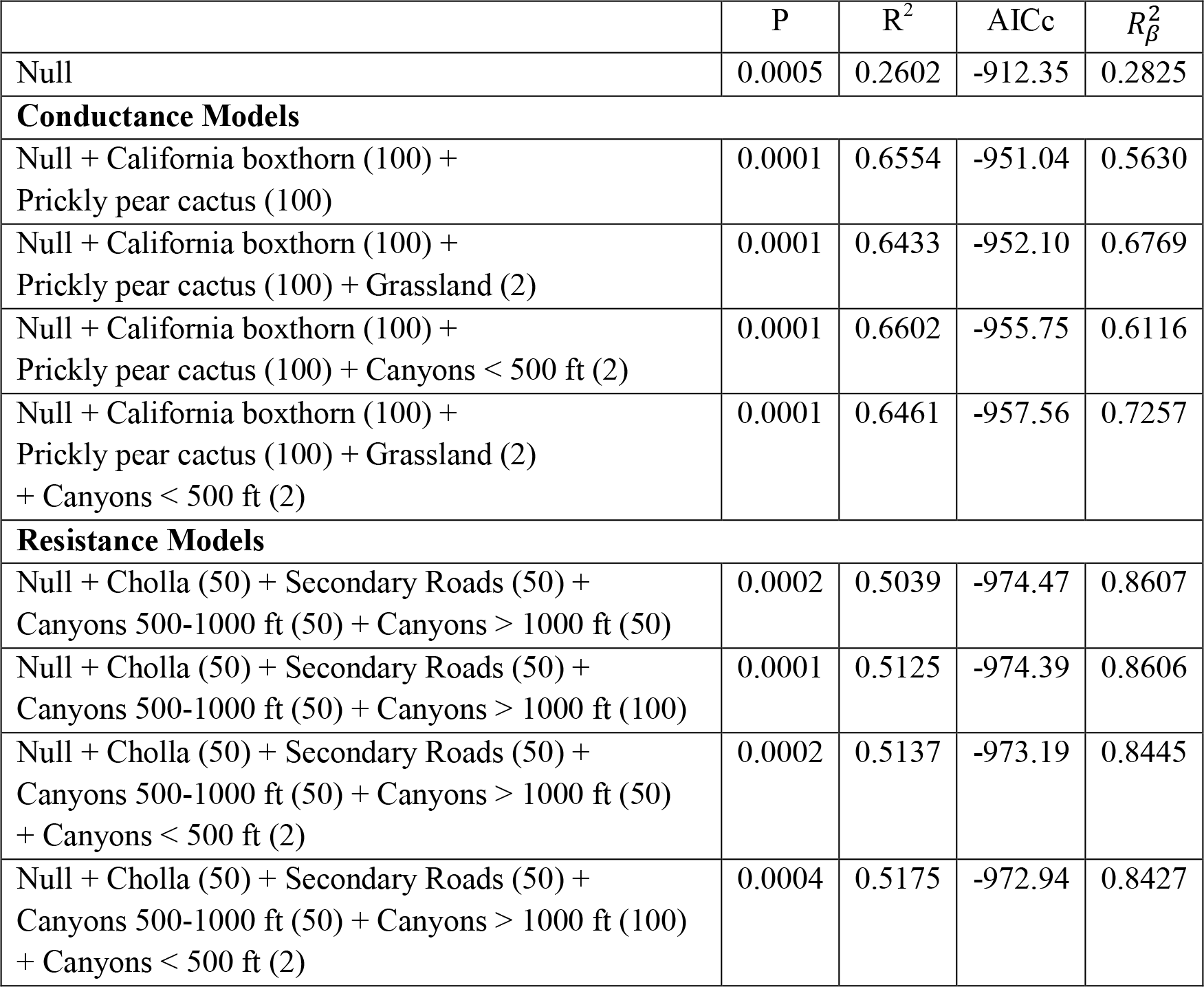
San Clemente Island competing conductance and resistance models. The best-supported conductance and resistance models are shown with the associated p-value (p) and coefficient of determination from non-parametric rank-based regressions (R^2^) from the resulting MRDM analysis. The corrected AIC value (AlCc) and 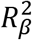 value are also presented and were used in model selection. Model components are listed with the value assigned to each factor in parentheses.

| | P | $R^2$ | AICc | $R^2_{\beta}$ |
| --- | --- | --- | --- | --- |
| Null | 0.0005 | 0.2602 | -912.35 | 0.2825 |
| **Conductance Models** |  |  |  |  |
| Null + California boxthorn (100) + Prickly pear cactus (100) | 0.0001 | 0.6554 | -951.04 | 0.5630 |
| Null + California boxthorn (100) + Prickly pear cactus (100) + Grassland (2) | 0.0001 | 0.6433 | -952.10 | 0.6769 |
| Null + California boxthorn (100) + Prickly pear cactus (100) + Canyons < 500 ft (2) | 0.0001 | 0.6602 | -955.75 | 0.6116 |
| Null + California boxthorn (100) + Prickly pear cactus (100) + Grassland (2) + Canyons < 500 ft (2) | 0.0001 | 0.6461 | -957.56 | 0.7257 |
| **Resistance Models** |  |  |  |  |
| Null + Cholla (50) + Secondary Roads (50) + Canyons 500-1000 ft (50) + Canyons > 1000 ft (50) | 0.0002 | 0.5039 | -974.47 | 0.8607 |
| Null + Cholla (50) + Secondary Roads (50) + Canyons 500-1000 ft (50) + Canyons > 1000 ft (100) | 0.0001 | 0.5125 | -974.39 | 0.8606 |
| Null + Cholla (50) + Secondary Roads (50) + Canyons 500-1000 ft (50) + Canyons > 1000 ft (50) + Canyons < 500 ft (2) | 0.0002 | 0.5137 | -973.19 | 0.8445 |
| Null + Cholla (50) + Secondary Roads (50) + Canyons 500-1000 ft (50) + Canyons > 1000 ft (100) + Canyons < 500 ft (2) | 0.0004 | 0.5175 | -972.94 | 0.8427 |

**Table 3.**
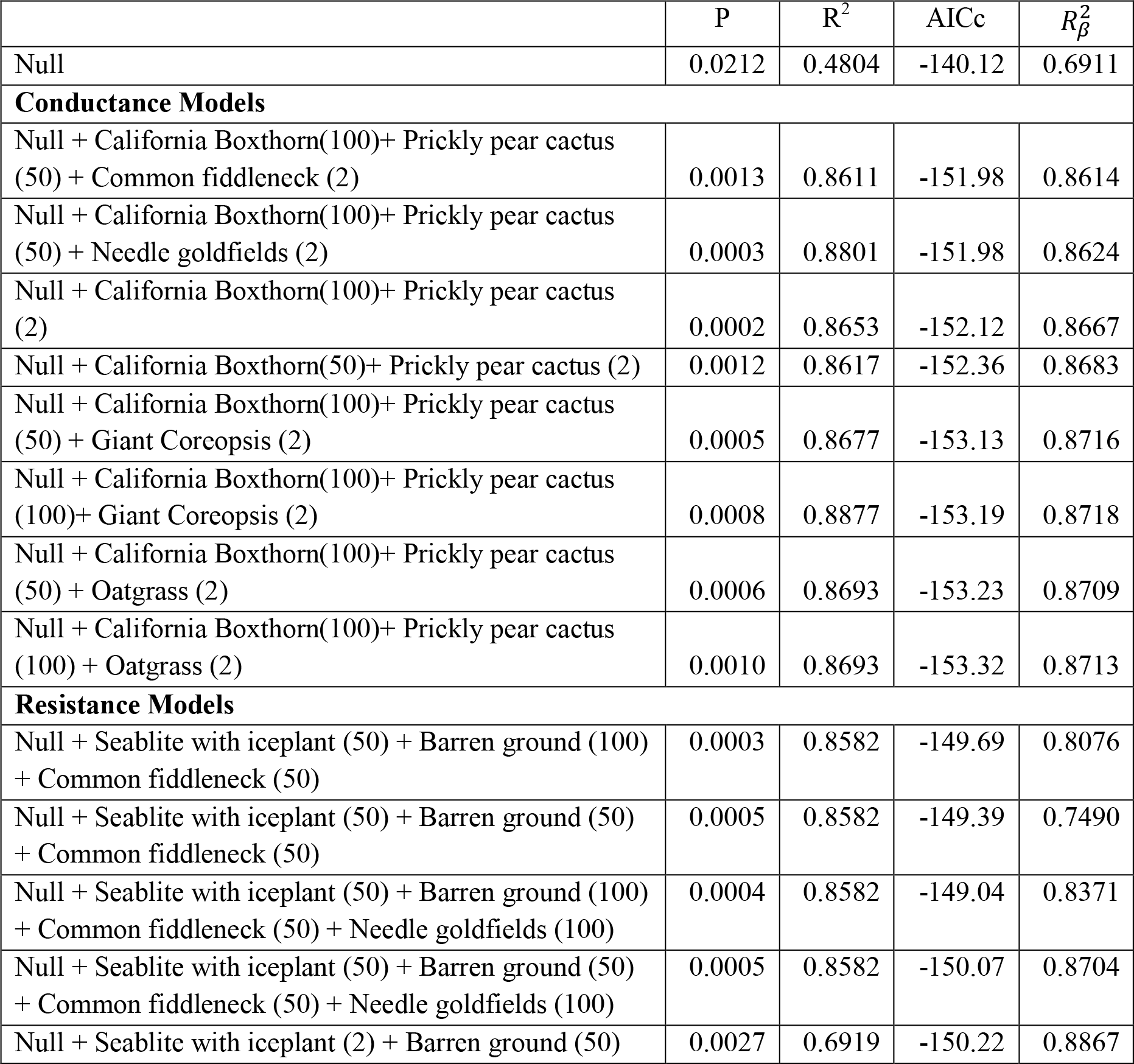
Santa Barbara Island competing conductance and resistance models. The best-supported conductance and resistance models are shown with the associated p-value (p) and coefficient of determination from non-parametric rank-based regressions (R^2^) from the resulting MRDM analysis. The corrected AIC value (AICc) and 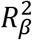 value are also presented and were used in model selection. Model components are listed with the value assigned to each factor in parentheses.

## Discussion

We used a single marker set to conduct a detailed genetic assessment of *X. riversiana* on San Clemente Island and Santa Barbara Island to inform conservation and management approaches across these independently managed islands. We used island-wide sampling with 23 microsatellite loci to estimate baseline parameters for each collection site, quantify population structure, and identify landscape factors correlated with genetic distance on each island. We found island night lizards were admixed on each island with shallow, but significant, population structure. Each island also shared common patterns of isolation by distance, with California boxthorn and prickly pear cactus identified as the most conductive habitat. Active sand dunes were identified as a barrier to gene flow on San Clemente Island but are absent on Santa Barbara Island. Unlike conductive habitat, habitat resistance patterns were unique to each island and included both natural and anthropogenic features.

### Population Structure

Structure identified 2 populations on San Clemente Island associated with the northern and southern ends and with strong admixture across the island (Figure 2). Surprisingly, Structure identified 4 populations on Santa Barbara Island despite its much smaller size. These populations were strongly associated with the western collection site (WP), northern collection site (AP), mid-island collection sites, and the south-eastern collection sites (Figure 3). The western and northern populations on Santa Barbara Island displayed relatively little admixture whereas other populations were admixed.

It is notable that the global Fst estimates for San Clemente Island and Santa Barbara Island were remarkably similar (0.0341 and 0.0346 respectively) despite large differences in island size and sampling scale. Our detection of population genetic structure on each island could result from the clustered sampling of a ubiquitously distributed organism characterized by limited dispersal (Schwartz and McKelvey 2009). However, congruence between methods corroborates the finding that multiple genepools exist across each island even at distances less than 400 m.

### Factors Positively Affecting Connectivity

California boxthorn and prickly pear cactus support the greatest abundances of island night lizards and are prime habitat types subject to continued monitoring (USFWS 2014). Our analyses identified prime habitat as highly conductive on both islands, which is supported by ecological data (Fellers and Drost 1991; Mautz 1993). Surprisingly, grasslands were identified as conductive on San Clemente Island even though previous research indicated that grasslands are poor habitat (Fellers and Drost 1991). Grasslands and small canyons on San Clemente Island may be conductive due to the presence of foraging resources and cover items. Cover items are important landscape features for thermoregulation (Regal 1968; Fellers and Drost 1991) and any habitat with suitable cover and resources may be conductive. For example, island night lizards on San Nicolas Island can be found in beach habitat with little vegetation, so long as cover rocks are present (Fellers et al. 1998). The Santa Barbara Island conductance model also included giant coreopsis as weakly conductive. The distribution of this habitat type on Santa Barbara Island is generally contiguous with prime habitat and difficult to analyze independently.

### Factors Negatively Affecting Connectivity

No habitat features were identified as resistant on both San Clemente Island and Santa Barbara Island. This finding probably reflects differences in topography, the extent of certain vegetation types, and history of anthropogenic development. Each island is thus characterized by unique factors which correlate with increased effective distances between collection sites and may constrain dispersal and connectivity. Active sand dunes on San Clemente Island were the only habitat we identified as an absolute barrier to dispersal, which is supported by Mautz (1993). There are no sand dunes present on Santa Barbara Island. It is likely that active sand dunes on San Nicolas Island act as barriers as well, although this hypothesis has yet to be assessed.

On San Clemente Island, cholla cactus may act as a resistant surface as the spines readily pierce flesh and could injure or debilitate fleeing or dispersing individuals (personal obs.). The southern bombardment area of San Clemente Island is of particular interest as increased disturbance may further propagate cholla increasing its density and distribution. If further fragmentation of populations within the southern portion of San Clemente Island becomes a management concern, we recommend controlling or minimizing the propagation of cholla through habitat restoration or more targeted training activities. Cholla habitat occurs in sparse and small patches on Santa Barbara Island likely resulting in the lack of congruent detection across islands.

On Santa Barbara Island, multiple habitat types were identified as resistant: common fiddleneck, needle goldfields, barren ground, and seablite grouped with iceplant. The identification of common fiddleneck and needle goldfields within similarly supported resistance models are not consistent with the ecological study of Fellers and Drost (1991) and could represent biases in the statistical optimization process. Notably, grassland habitats were absent from the final resistance model despite Fellers and Drost (1991) finding that grasslands required equivalent capture efforts to seablite and iceplant habitats. It is likely that when cover is dense enough to cause refugial fissures in the soil or supplemented by cover items, resistance may be minimized. Iceplant may render areas unsuitable for other vegetation (Vivrette and Muller 1977) and may, as with barren ground, limit the cover necessary for foraging, thermoregulation, and dispersal (Fellers and Drost 1991). However, we could not determine the independent contributions of seablite and iceplant habitats due to the spatial and temporal transitions between them (pers. comm., Rodriguez 2016). Thus, on Santa Barbara Island, we recommend habitat remediation efforts which prioritize iceplant removal and revegetation with prime habitat.

Canyons >500 ft in length and secondary roadways were also detected as resistant on San Clemente Island. The association of canyons with increased genetic differentiation between collection sites is supported by ecological research, and may be due to both difficulty in crossing canyons and a lack of resources within them (Mautz 1993). Secondary roadways form extensive networks which fragment prime habitat of boxthorn and prickly pear with barren ground, particularly along the western escarpment of the island. Notably, the primary roadway was not identified as a resistant surface, even though it is paved and has more traffic. However, this primary road is a relatively short road running down the middle of the island and is unlikely to significantly influence connectivity patterns. Thus, any efforts to mitigate fragmentation should be applied to secondary roadways.

### Management Implications

Management of species that persist on oceanic islands needs to examine the conductance and resistance of habitat within islands and compare between them to derive generalizable actions. As many management actions are spatially explicit, it is important to characterize genetic patterns using spatially explicit methods (Shaffer et al. 2015). The workflow used in our study can be generalized to examine connectivity patterns within mainland ecosystems that are fragmented into habitat islands to inform management actions within and between independently managed habitat fragments.

An exemplar region which could benefit from this approach is the peninsular region of Southern California, which is separated into several administrative units tasked with management and monitoring activities. This region may experience increased extinction risks of plant and animal species (e.g. Lawson 2011) and genetic studies of terrestrial vertebrates have revealed patterns of significant genetic differentiation and decreased genetic diversity (Luckau 2015; Lion 2016). The modeling frameworks we present can be applied to support management efforts of peninsular species through the identification of fine-scale resistant features, for targeted mitigation efforts within each management area, while also identifying key landscape features within and between management areas which support connectivity. This combined approach may better identify management strategies aimed at sustaining local populations, such as increasing the area of prime habitat and improving the quality of the intervening matrix (e.g. Fahrig 2001).

The contemporary genetic patterns we identify for *X. riversiana* can inform management and monitoring practices on each independently managed island. Common management strategies across islands include restoring invasive vegetation to prime habitat to support intra-island connectivity and increase population sizes. Improving intra-island connectivity and population size may also increase species resilience to potential threats of climate change or other catastrophic events by increasing genetic diversity and decreasing the effects of demographic stochasticity. We encourage the comparison of fine-scale patterns within and between regional management units to identify common strategies and unique factors to inform mitigation efforts.

## Acknowledgements

All capture data, individual genetic profiles, and protocols used to generate microsatellite data may be found in SR’s dissertation (Rice 2017). We would like to thank the National Park Service for access to Santa Barbara Island and providing landscape-level GIS layers, San Clemente Naval Base for access to San Clemente Island and providing landscape-level GIS layers, and our field assistants. In addition we would like to thank Drs. A. Bohonak, H. Regan, K. Anderson, J. Gatesy, and R. Rose for comments received during manuscript preparation.

